# Examining the efficacy of localised gemcitabine therapy for the treatment of pancreatic cancer using a hybrid agent-based model

**DOI:** 10.1101/2022.04.18.488716

**Authors:** Adrianne L. Jenner, Wayne Kelly, Michael Dallaston, Robyn Araujo, Isobelle Parfitt, Dominic Steinitz, Pantea Pooladvand, Peter S. Kim, Samantha J. Wade, Kara L. Vine

**Affiliations:** School of Mathematical Sciences, Queensland University of Technology, Brisbane, QLD, Australia; School of Computer Science, Queensland University of Technology, Brisbane, QLD, Australia; Tweag Software Innovation Lab, London, United Kingdom; Kingston University, Kingston, United Kingdom; School of Mathematics and Statistics, University of Sydney, NSW, Australia; Illawarra Health and Medical Research Institute, Wollongong, NSW, Australia; School of Chemistry and Molecular Bioscience, University of Wollongong, Wollongong, NSW, Australia

## Abstract

The prognosis for pancreatic ductal adenocarcinoma (PDAC) patients has not significantly improved in the past 3 decades, highlighting the need for more effective treatment approaches. Poor patient outcomes and lack of response to therapy can be attributed, in part, to the dense, fibrotic nature of PDAC tumours, which impedes the uptake of systemically administered drugs. Wet-spun alginate fibres loaded with the chemotherapeutic agent gemcitabine have been developed as a potential tool for overcoming the physical and biological barriers presented by the PDAC tumour microenvironment and deliver high concentrations of drug to the tumour directly over an extended period of time. While exciting, the practicality, safety, and effectiveness of these devices in a clinical setting requires further investigation. Furthermore, an in-depth assessment of the drug-release rate from these devices needs to be undertaken to determine whether an optimal release profile exists. Using a hybrid computational model (agent-based model and partial differential equation system), we developed a simulation of pancreatic tumour growth and response to treatment with gemcitabine loaded alginate fibres. The model was calibrated using *in vitro* and *in vivo* data and simulated using a finite volume method discretization. We then used the model to compare different intratumoural implantation protocols and gemcitabine-release rates. In our model, the primary driver of pancreatic tumour growth was the rate of tumour cell division and degree of extracellular matrix deposition. We were able to demonstrate that intratumoural placement of gemcitabine loaded fibres was more effective than peritumoural placement. Additionally, we found that an exponential gemcitabine release rate would improve the tumour response to fibres placed peritumourally. Altogether, the model developed here is a tool that can be used to investigate other drug delivery devices to improve the arsenal of treatments available for PDAC and other difficult-to-treat cancers in the future.

**Author Summary:** Pancreatic cancer has a dismal prognosis with a median survival of 3-5 months for untreated disease. The treatment of pancreatic cancer is challenging due to the dense nature of pancreatic tumours which impedes retention of drug at the tumour site. As such, systemic administration of chemotherapies, such as gemcitabine, has a limited efficacy. To overcome this, sustained-release devices have been proposed. These devices are injected locally and release drug slowly over time, providing a concentrated local, sustained drug concentration. To investigate the possible efficacy of these devices, we developed a mathematical model that would allow us to probe treatment perturbations *in silico*. We modelled the individual cancer cells and their growth and death from gemcitabine loaded into the sustained delivery devices. Our platform allows future investigations for these devices to be run *in silico* so that we may better understand the forms of the drug release-profile that are necessary for optimal treatment.

## Introduction

Inoperable pancreatic ductal adenocarcinoma (PDAC) has a dismal prognosis, with a median survival of 3−5 months for untreated disease [1]. Treatment of PDAC with the chemotherapeutic agent gemcitabine can achieve clinical benefit and symptom improvement in 20−30% of patients [1, 2], although PDAC is still regarded as a chemotherapy-resistant tumour [3, 4]. Gemcitabine is designed to target and kill cancer cells by incorporating into the DNA strand of a PDAC cell allowing only one deoxynucleotide to be incorporated, which prevents strand elongation [5, 6], resulting in cell cycle arrest and subsequent cell death [7, 8]. Despite gemcitabine being established as a standard treatment for advanced PDAC over 20 years, most subsequent large phase III studies have not shown significantly improved survival benefit [9]. Overall prognosis for PDAC has seen little improvement in the last 3 decades, largely due to drug resistance and poor intratumoural drug accumulation.

The majority of chemotherapeutics, gemcitabine included, are administered systemically via bolus or infusion intravenous administration. This often results in significant systemic toxicity, with only a fraction of the injected dose reaching the tumour. As such, there has been a growing interest in the development of localized targeted delivery systems which can modify the bio-distribution of drugs and achieve local drug accumulation in the tumour tissue [10–12] (**Figure 1**). For example, drug-eluting polymeric implants are designed to deliver high concentrations of chemotherapeutic drugs directly at the tumour site, overcoming transport and tissues barriers as well as limiting off-target toxicities [13]. Biodegradable implants, can be designed to provide sustained drug release over weeks or months, avoiding repeated external drug dosing, clinic visits and other surgical interventions. The characteristics of these devices make local delivery especially attractive for chemotherapeutics with a narrow therapeutic window or short *in vivo* half-life [14], such as gemcitabine.

Drug-loaded polymeric fibres can be prepared by various cross-linking methods and allow for drug molecules to be released in a controlled manner depending on the cross-linking type and methods [15]. Previously, Wade *et al*. [14] showed that wet-spun gemcitabine-loaded alginate fibres inhibited *ex vivo* PDAC spheroid growth and reduced PDAC cell viability compared to gemcitabine delivered as a free drug. In a subsequent study, Wade *et al*. [13, 16] showed that a coaxial fibre formulation, in which the alginate was encased by a polycaprolactone (PCL) shell demonstrated significant *in vivo* antitumour efficacy; however, it is not possible to conclude experimentally whether an alternative release-profile of gemcitabine may be more effective. Fortunately, computational and mathematical modelling is well situated as a predictive tool for quantifying the efficacy of alternative drug release profiles and drug administration patterns.

Mathematical models have been used to help understand formation and treatment of a range of different cancers for some time now [17–20]. In particular, agent-based models (ABM) have been used extensively in cancer modelling as they allow for the consideration of spatial and phenotypic heterogeneity [21–27] which are known to be major drivers of variations in treatment outcomes. In ABMs, the likelihood of events, such as cell proliferation, movement, death or mutation are modelled as probabilities, allowing the simulation to evolve stochastically in time. Phillips *et al*. [28] presented a hybrid mathematical approach that characterized vascular changes during tumour growth via an ABM, with treatment, nutrient, and VEGF changes captured through a continuum model. Insights on therapeutic failure in immunotherapy have also been provided through an ABM software known as PhysiCell [29, 30]. Oncolytic virotherapy has also been the focus of numerous ABMs [31–35], with an ABM of virotherapy demonstrating that the parameter range leading to tumour eradication is small and hard to achieve in 3D. There have been ABMs developed that specifically focus on pancreatic cancer growth [36, 37]; however, an ABM describing pancreatic cancer growth and treatment with a degradable polymer implant has not yet been developed.

For some time, mathematical models of degradable drug delivery mechanisms have been used to assist in the understanding of polymer degradation, hydrolosis kinetics and the subsequent effect of drug release on the applied system [10, 38–45]. Using mass-balance kinetic equations, McGinty *et al*. [42] investigated the extent to which variable porosity drug-eluting coatings can provide better control over drug release using transport diffusion equations. Their results indicate that the contrast in properties of two layers can be used as a means of better controlling the release, and that the quantity of drug delivered in early stages can be modulated by varying the initial drug distribution.

More recently, Spiridonova *et al*. [46] fitted drug release from polymer microparticles and investigated the effect of size distribution on diffusional drug release from sustained-delivery systems using a system of partial differential equations (PDEs). Whilst useful for capturing the drug delivery mechanism, most models of drug-loaded polymers such as these have not examined the influence of changes to drug release profiles on antitumour efficacy or how intratumoural stochasticity impacts drug delivery.

In this work, we have developed a hybrid mathematical and computational model of PDAC tumour growth and death from treatment with gemcitabine released from a polymeric fibre. We extended a previously published ABM known as a Voronoi cell-based model (VCBM) [32] to model tumour cell growth and death and coupled this with a PDE model for gemcitabine release from polymeric implants. *In vitro* drug release curves were used to optimise the PDE formulation describing how gemcitabine is released from fibres. A numerical simulation was then used to initialise the parameters in the ABM using *in vivo* control PDAC tumour growth measurements. The potential impact of these fibres on tumour growth and cell death was then investigated with the VCBM-PDE model and improvements on drug release kinetics and fibre placement were suggested. The model was developed as a tool that can be applied to interrogate other cancer therapies using polymeric implants with the goal to improve treatment response for PDAC patients.

## Experimental methods

### Fibre fabrication and characterisation

Full details for the fabrication and characterisation of alginate fibres loaded with our without gemcitabine are described in Wade *et al*. [13, 14]. Briefly, gemcitabine-loaded alginate fibres had a uniform surface area from 50 − 120 µm in diameter. Fibres displayed different drug release profiles depending on the concentration of polymer used. Fibre diameter also varied depending on the materials used [14].

### Fibre gemcitabine release kinetics

Full details for the experiments measuring gemcitabine release can be found in Wade *et al*. [14] with brief details here. Gemcitabine-loaded fibres were added to 2mL of simulated body fluid (SBF), Ph 7.4 and incubated at 37°C. At various time points (10, 30, 60, 90 min hourly for 10h and then daily for 3 weeks), buffer solution (200µL) was removed for analysis of gemcitabine release and replaced with fresh SBF. The amount of drug released from alignate fibres was determined using high performance liquid chromatography (HPLC). The amount of gemcitabine released (µg) was calculated by interpolating AUC values from the standard curve using Empower Pro V2 (Waters) software.

### Implant toxicity in vitro

Gemcitabine loaded fibres were tested for their cytotoxicity against human pancreatic cancer cells (Mia-PaCa-2) cells over 72h. Cells were incubated with 0.5 cm lengths of gemcitabine loaded or non-drug loaded fibre formulation before an endpoint MTS cell viability assay was performed. Results are displayed as a percentage of an untreated control. Experiments were performed in triplicate. Full details for the toxicity experiments can be found in Wade *et al*. [13].

### In vivo Mia-PaCa-2 cell growth

Animals were subcutaneously inoculated with 100µL suspension of 1 × 10^6^ Mia-PaCa-2 cells in PBS. Tumour volume measurements began when tumours reached a volume of 200 *mm*^3^ using

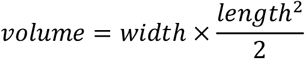

where *w* is the longest tumour measurement and *l* is the tumour measurement along a perpendicular axis. Tumour volume was measured daily for a duration of approximately 33 days. Full details for this experiment can be found in Wade *et al*. [13]. All animal experiments were conducted in accordance with the NHMRC Australian Code for the Care and Use of Animals for Scientific Purposes, which requires 3R compliance (replacement, reduction, and refinement) at all stages of animal care and use, and the approval of the Animal Ethics Committee of the University of Wollongong (Australia) under protocol AE18/13.

## Mathematical methods

The model developed for the release of gemcitabine from alginate fibres and the impact on a growing PDAC tumour was formulated in two parts. The first describes the PDE describing the concentration of gemcitabine in the tumour microenvironment (TME) and surrounding tissue over time. The second describes the VCBM [32] that captures the way tumour cells proliferate, move and undergo apoptosis from gemcitabine. All parameters introduced for the model are summarised in **Table S1-S5** in the **Supplementary Tables and Figures** and a schematic for the model is in **Figure 2**.

### Model of gemcitabine

To capture the concentration of gemcitabine in the tumour microenvironment, we first considered a 2D rectangular domain with boundary *B* (**Figure TS1**). Inside this domain, is implanted a gemcitabine drug-loaded fibre which is represented by a vertical line source (**Figure TS2**A and **Figure 2**A). Gemcitabine diffuses from the line source at some time-dependent rate that decreases as the polymeric fibre degradation slows. The gemcitabine concentration in the domain is diffusing and decaying. PDAC cells in the domain are also taking up gemcitabine, removing it from the concentration in the domain. Inside the fibre, we model the diffusion of drug as radially symmetric (**Figure 2**B).

**Figure 1.**
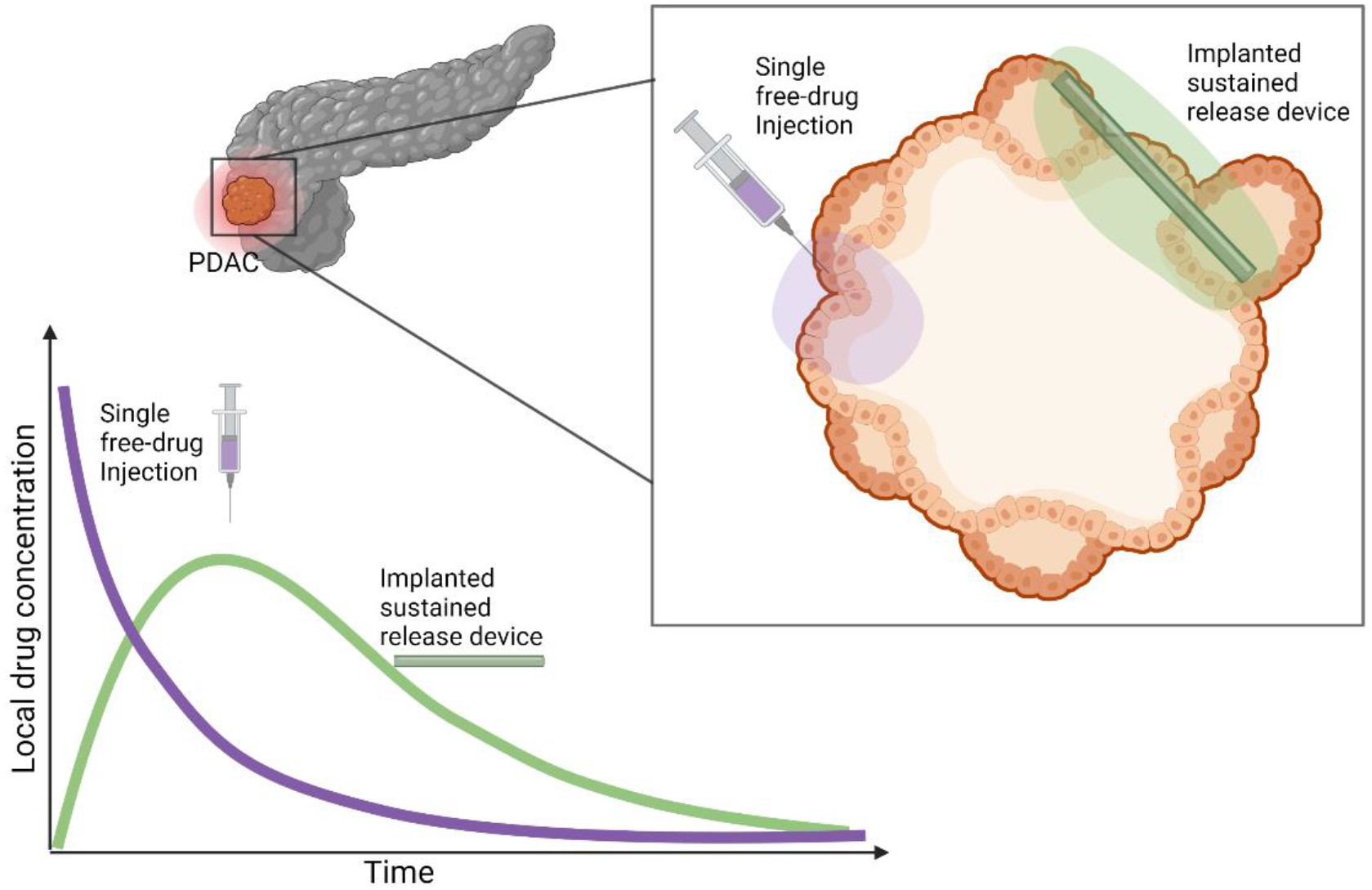
Motivation for sustained-delivery implants for treatment of PDAC. Sustained-delivery implants are a promising treatment methodology over conventional single free-drug intravenous or intrantumoural injections. A hypothetical comparison of drug concentrations at the tumour site under these two protocols is pictured. Systemic injections of anti-cancer drugs often result in a rapid decrease of drug concentration at the tumour site. In comparison, sustained-release mechanisms deliver drug over a prolonged period resulting in a durable drug presence at the tumour site. Created using biorender.com.

We denote the concentration of drug in the TME at position (*x, y*) by *C*(*x, y, t*) and model this concentration by

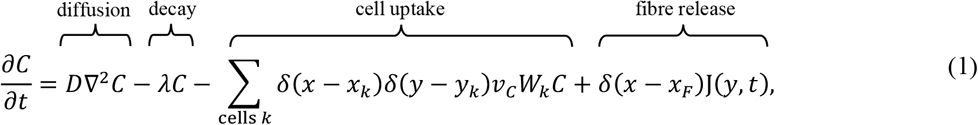

where *D* is the diffusion coefficient in the TME, and λ is the decay rate of the drug. To model cancer cells taking up gemcitabine, we used *δ*(*x*) which is the Dirac delta function in one-dimension, where (*x*_*k*_, *y*_*k*_) is the *k*th cancer cell’s Voronoi centre position in the domain (**Figure S1**), and *W*_*k*_ is the cell’s volume. Pancreatic cancer cells take up drug in the domain at a rate *v*_*c*_. Cell uptake was modelled by point sinks analogous to that in PhysiCell and BioFVM [30, 47], where cells are considered discrete “point masses” in the domain that take up drug from a single rectangular discretized voxel weighted by the local concentration of drug. We then used a line source at *x* = *x*_*F*_, *y*_0_ ≤ *y* ≤ *y*_0_ + *L* to model the release of gemcitabine from the polymeric fibre, where *y*_0_ is the location of the bottom of the fibre and *L* is the fibre length (**Figure TS1**). This line source was represented by a Dirac delta function in one-dimension and the drug diffuses from the line source with flux J(*y, t*).

To derive the flux of drug from the line source, we first assumed that the release of drug from the fibre would be time dependent. As such, we chose to explicitly model a concentration of drug diffusing inside the fibre. We denote the concentration of gemcitabine at radial position *r* and location (*x*_*F*_, *y*) by *F*(*r, y, t*) (**Figure TS2** and **Figure 2**A). We model the diffusion and movement of drug inside the fibre assuming radial symmetry. We assumed that diffusion in the radial direction is significantly faster than along the fibre since the radius of the fibre *r*_*total*_ is significantly less than the length of the fibre *L* (**Figure TS1** and **Figure TS2**). This gives

**Figure 2.**
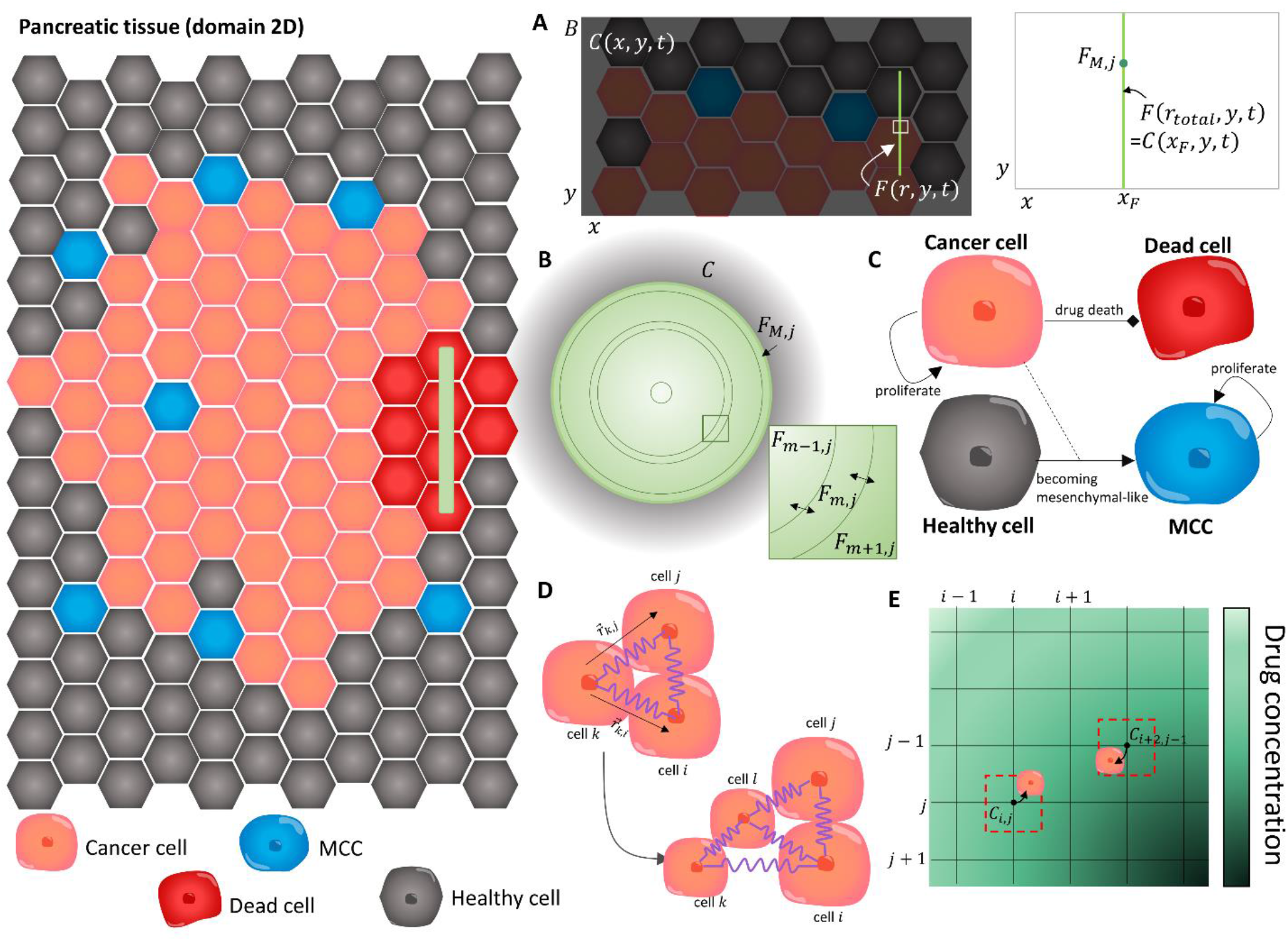
The main components of the VCBM-PDE model. (A) The concentration of drug in the TME was modelled in a 2D domain bounded by *B*, where *C*(*x, y, t*) was the concentration in the TME at position (*x, y*). The fibre implant was then placed at a position *x* = *x*_*F*_ and modelled as a line source. To capture the diffusion of drug from the fibre, we modelled the concentration of gemctiabine inside the fibre *F*(*r, y, t*) at radial position *r* and domain position *y* where the continuity condition in **Eq. (3)** required equal concentrations at the fibre boundary and at the immedicate local microenvironment, i.e. *F*(*r*_*total*_, *y, t*) = *C*(*x*_*F*_, *y, t*). (B) The concentration of gemcitabine inside the polymeric fibres was modelled by radially symmetric difussion **Eq. (2)** using a finite volume method (FVM) discretisation and considering the 2D cylindrical cross section of the fibres which have length *L* and radius *r*_*total*_. The fibre was discretised into concentric annuli *F*_*m,j*_ at annulus *m* andcross section *j*, (*i* = 0,1, …, *M*) and the concentration of drug in each annulus *F*_*m,j*_ was modelled by considering drug diffusion across the bounadaries (e.g. *F*_*m*−1,*j*_ and *F*_*m*+1,*j*_ flow into *F*_*m,j*_ and vice vera). The full discretisation is presented in the **Technical Supplementary Information**). (C) Modelling assumptions for the VCBM were that cancer cells (pink) proliferate and some are able to cause epithelail to mesenchymal transtion and become invasive. We model this transition by assuming cells differentiate into an mesenchymal cancer cell (MCC) with one daughter cell placed on a neighbouring healthy cell. These MCCs cause the break down of surrounding tissue (i.e. replace healthy neighbouring cells with their progeny). Cancer cells can then die through gemcitabine uptake from their local environment. (D) Individual cells were modelled as cell centres connected by springs [32]. The proliferation of a cell introduced a new cell into the lattice network which caused the rearrangement of the cells in the lattice with movement governed by Hooke’s law. (E) To simulate the gemcitabine concentration in the TME, **Eq. (1)**, we introduced a FVM discretisation, where the gemcitabine concentration was defined at discrete volumes centered around points in the discretisation. Cells could take up drug from the nearest grid point to their centre, and this concentration was used to determine their likelihood of drug-induced cell-death.

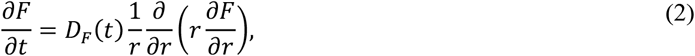

where *D*_*F*_(*t*) is the time-dependent diffusion of drug inside the fibre. We imposed the continuity condition

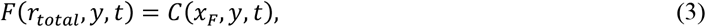

so that the diffusion of drug out of the fibre at the line source will depend on the location (*x*_*F*_, *y*) and local exterior concentration. The flux out of the line source J(*y, t*) in **Eq. (1)** can then be approximated from the release of drug across the boundary of the fibre:

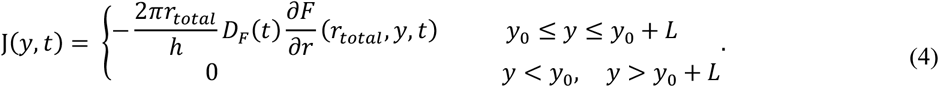

This term is derived by converting the flux out of the radial fibre into the flux represented by the line source in **Eq. (1)** and converting to a concentration per surface area where *h* is the depth of the rectangular region (presumed thing, see **Figure TS1**). Both **Eq. (3)** and **Eq. (4)** are necessary boundary conditions for **Eq. (1)** and **Eq. (2)**. In this way, we assume the concentration is continuous and the flux of the fibre is equal to the flux into the TME, equivalent to a conservation of mass.

The diffusivity of the drug, *D*_*F*_(*t*), is modeled by the function

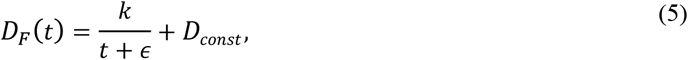

where *k* controls the decay rate to the constant decay rate from the fibre (i.e. how quickly the fibre swells), *D*_*const*_ is the constant decay rate from the fibre and *ϵ* is a tuning constant to provide a finite initial diffusion coefficient, i.e. *D*_*F*_(0) = *k*/*ϵ* + *D*_*const*_. We expect *D*_*F*_(0) to be initially large (>1) since the polymeric fibre is hydrophilic and drug would immediately diffuse out of the fibre. In addition, some drug is never properly loaded into the fibre and can be released instantaneously. The formalism in **Eq. (5)** was broadly chosen to capture the rapid decline in release as the polymeric fibre degrades. It is possible to model the breakdown of the drug release mechanisms to include device swelling and degradation and for examples of this see [46, 48–50].

No-flux boundary conditions on *B*, the exterior of the TME, are imposed:

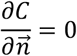

where 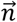 is the outward unit normal on the boundary *B* (**Figure TS5**). In the case of a fibre implantation, all drug in the domain is initially situated in the fibre:

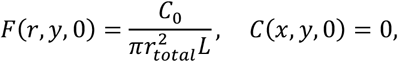

where *C*_0_ is the amount of drug in *µg*, the denominator is the volume of the fibre and there is no drug initially in the domain *B*. We assume the location of the fibre is fixed in space over the course of the simulation and is not affected by cells around it. For more details on the derivation of the model see the **Technical Supplementary Information**.

We solved **Eqs. (1)-(4)** numerically using a Finite Volume approximation. In particular, the diffusion of drug within the fibre, **Eq. (6)**, was solved through discretising the cross section of a fibre into annuli (see **Figure 2**B and the **Technical Supplementary Information**). The model is solved using a finite volume method (FVM) discretization, for examples of this form of discretization in cancer growth and treatment see [51–59].

### Voronoi Cell-Based Model (VCBM) of pancreatic tumour growth

Agent-based models (ABMs) are primarily used to simulate heterogeneity that arises through stochasticity in cellular interactions. We present an ABM to capture the 2D formation of a pancreatic tumour in the pancreas. Our model extends a Voronoi cell-based model (VCBM) for tumour growth already published in [32]. The model describes how individual cells behave over time by considering their behaviour to be a stochastic process. It uses points as representatives of cell centres and then overlays this with a Voronoi tessellation to define individual cell boundaries. A Voronoi tessellation defines the region of space where the Euclidean distance to a point is less than the distance to any other cell centre in the lattice. Voronoi tessellations have been used to model tissue and cancer cell dynamics for some time [60–64]. Using a Voronoi tessellation for the ABM allows cell morphology to be heterogeneous and not fixed, and the morphology can change with cell movement. The model is solved on a time increment of 1hr to account for the fact that cellular interactions are slow in comparison to drug diffusion (**Figure TS3**). To model pancreatic tumour formation, we assumed the primary functions of pancreatic tumour cells were movement and proliferation. Below are details of the cell types, the model for cell movement and proliferation, a description of the dynamics of tumour mesenchymal cells, the model for cell death and details of how the domain changes as the tumour grows.

PDAC cells can acquire mesenchymal-like phenotype properties through a process known as epithelial-mesenchymal transition (EMT) [65–68]. In the EMT process, epithelial elements undergo cytoskeleton remodelling and migratory capacity acquisition due to the loss of intracellular contacts and polarity [66]. This enables the formation of mesenchymal-like cancer cells (MCCs) which have enhanced migratory capacities and invasiveness, as well as elevated resistance to apoptosis [67]. Since there is evidence that EMT plays an important role in PDAC progression [65–68], we have introduced this cell type into the model.

We considered four main cell types in the model: healthy pancreatic cells, PDAC cells, MCCs and dead cells (cancer cells that have experienced drug-induced death), see **Figure 2**C. The initial tissue comprised of healthy cells, arranged so that the corresponding Voronoi cells form a hexagonal tessellation, analogous to other work in the literature [69, 70]. To initialise the tumour formation, we removed a healthy cell from the centre of the domain and replaced it with a pancreatic tumour cell (**Figure S1, Supplementary Tables and Figures**). These pancreatic tumour cells could proliferate, die from gemcitabine, or form MCCs. Once formed, these pancreatic stem cells then move and proliferate until they die. Healthy cells are assumed to be able to move or become MCCs.

Cell movement is governed by pressure-driven motility, modelled using Hooke’s law [32]. Each cell’s position is updated by calculating the effective displacement of the cell’s lattice point by the sum of the forces exerted on that cell, where force is modelled as a network of damped springs connecting a cell to its nearest neighbours (defined by a Delaunay triangulations). Consider cell *k*, the displacement of this point in time Δ*t*_*cells*_ is given by

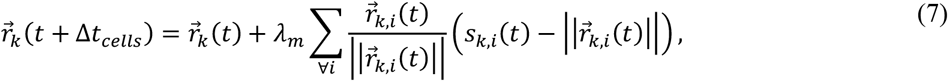

where 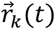 is the position of the *k*th point in the lattice at time *t*, λ_*m*_ is a damping and mobility constant, 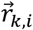 is the vector between *k* and *i, s*_*k,i*_ is the spring rest length (equilibrium distance) between cell *k* and *i*. The introduction of new cells in the lattice through proliferation introduces new spring connections and shortens or extends others, promoting the movement of cells in the environment (**Figure 2**D).

Tumour cell proliferation was assumed to be a function of the cell’s distance, *d*_*neut*_, to the nutrient source (tumour periphery, i.e. nearest healthy cell centre, see **Figure S3**). The maximum radial distance for nutrient-dependent cell proliferation is *d*_*max*_. Cells that are a further distance from the nutrients than *d*_*max*_ enter a quiescent (non-proliferative state), forming what is commonly known as a necrotic core. The probability of a cell dividing *p*_*d*_ in time step Δ*t*_*cells*_ is given by

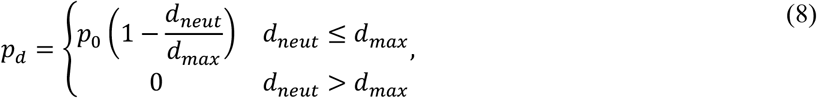

where *p*_0_ is a proliferation constant derived based on the maximum rate of cell proliferation *r* (i.e. *p*_0_ = 1 − exp(−*r*Δ*t*) ≈ *r*Δ*t*). The formalism in **Eq. (8)** is similar to what was used by Kansal *et al*. [71], Jiao and Tarquato [72] and Jenner *et al*. [32]. A cancer cell’s ability to proliferate was also based on whether there was enough local space for proliferation to occur. If a cell *k* proliferates, a new lattice point *l* is created and the two cells are placed at a distance *s*/*p*_*age*_ from the original proliferating cells position at a rotation *θ* ∼∈ *U*(0,2*π*] (**Figure 2**D). To simulate the enlargement and repositioning of the daughter cells, the resting spring length of the connection between *k* and *l* linearly increases over time from *s*/*p*_*age*_ to the mature resting spring length *s* as was formulated in our previous work [32]. Once a cell has proliferated, it takes *g*_*age*_ time steps before the daughter cell will try to proliferate again, accounting for G1 phase of the cell cycle where the cell transitions from mitosis M to DNA synthesis S [32]. It is well known that tumours contain highly heterogeneous populations of cells that have distinct reproductive abilities. To account for heterogeneity in the cell cycling, cells sampled the age at birth from a Poisson distribution with mean 50.

MCCs are created at the boundary of the tumour with probability *p*_*MCC*_. These cells are created from tumour cells differentiation into a tumour cell and an MCC. We mode their invasive property by placing the daughter cell at the position of a neighbouring healthy cell, removing it from the domain. Through their creation, these MCCs contribute to the degradation of the healthy tissue surrounding the tumour.

As in [73–76], we assumed that cancer cells die from gemcitabine contact at a rate described by the Michaelis-Menten term

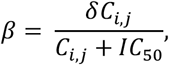

where *δ* is the maximum death rate due to the drug, *C*_*i,j*_ is the concentration of drug at the grid position (*i, j*) in the FVM discretization closest to the cell’s centre (**Figure 2**E and the **Technical Supplementary Information**), and *IC*_50_ is the concentration at which half the effect of the drug is attained. From this, the probability of an individual cell dying can be determined by assuming Prob(cell death)= 1 − exp(−*β*Δ*t*) ≈ *β*Δ*t*. While we chose not to model explicitly the resistance to gemcitabine that cancer cells can develop [3, 4], we believe that by modelling cell death probabilistically we can capture some of the heterogeneity that may exist intratumourally. If a cell dies, then its phenotype changes to be a dead cell and and takes *d*_*age*_ hours to disintegrate. To simulate disintegration, at each time increment the spring rest lengths of a dead cell to each of its neighbours, *s*_*k,i*_, decreases by *s*_*k,i*_/*d*_*age*_.

As the tumour grows, the model domain expands. To reduce computational cost, new healthy cells are added to the domain only when a tumour cell’s radial distance from empty space is < 10µ*m* (**Figure S2 Supplementary Tables and Figures**).

### Numerical simulations and parameter estimation

The VCBM-PDE model was written in C++ and simulations called through Matlab 2021b by creating a definition file for the C++ library using *clibgen* and *build* in Matlab 2021b. Code for the model at the various stages (e.g. fibre, single injections) can be found on github (https://github.com/AdrianneJennerQUT/hybrid-VCBM-of-gemcitabine-and-pancreatic-cancer). Full details on all aspects of the code can be found in **Code Documentation**.

An approximation for tumour volume was then determined from the 2D simulations using the same formula as the calibre measurements, multiplied by a scalar *σ*:

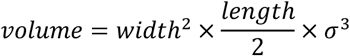

where *width* is the longest distance of a cell on the periphery from the centre and *length* is the distance of the farthest cell from the centre on the radial axis perpendicular to the radial axis of the longest distance (**Figure S5 Supplementary Tables and Figures**) where *σ* unit length of the model is equivalent to 1 mm. This calculation choice was made to closely resemble the tumour volume calculation with calibres done *in vivo*. As the size of the computational domain was smaller than the size of the real tumour, the length unit was scaled by *σ*, which scaled the unit length in the VCBM domain to a comparable mm unit measurement that reduced the computational cost. We chose *σ* = 0.1728.

All fitting was undertaken using *lsqnonlin* in Matlab 2021b using *pdepe* and *ode45* to simulate the model. Parameters in the model were fit using experimental data or estimated from the literature. To fit the parameters relating to drug release from the fibre we used the *in vitro* drug release experiments. We simplified the model to consider only one cross section, i.e. *F*_*m,j*_ = *F*_*m*_, since the outside concentration of drug was independent of location in the absence of cells in the *in vitro* experiment.

To estimate parameters for the pancreatic cell growth kinetics, we did a large Latin Hypercube sample of the parameter space and determined parameters that resulted in a minimal least squares distance to the *in vivo* control tumour growth measurements. Other parameters were either fixed to previous values in the literature or estimated based on previous work. See **Tables S1-S5** in **Supplementary Tables and Figures** for a full summary of all parameter values and relevant references.

## Results

### Calibration of drug release kinetics and drug-induced cell death to in vitro measurements

Gemcitabine-loaded fibres were placed in a solution bath and the resulting cumulative concentration of gemcitabine measured (**Figure 3**A). To obtain a model describing the release rate of the drug from the fibre, we fitted parameters from **Eq. (1)-(4)** to these *in vitro* measurements for the release of gemcitabine from 3% alginate 15% PCL fibres [14]. Fitting the release curve parameters *k, d*_*const*_, *C*_0_ and *A*_*out*_ gave the fit in **Figure 3**B and parameter values in **Table S1**. Overall, the model was able to obtain the fit to the data and followed the trend which showed a rapid initial release of gemcitabine followed by a steady-state threshold. We validated the model’s predictive capability by also fitting gemcitabine release from 1% and 2% alginate fibres (**Figure S4 Supplementary Tables and Figures**).

To assess the efficacy of the drug on inducing death in PDAC cells, cell viability studies were performed using Mia-PaCa-2 cell lines. To model these experiments, we considered a simplified deterministic and spatially independent version of our model with only live cancer cells *P*_*L*_ (*t*), dead cancer cells *P*_*D*_ (*t*) and a concentration of drug *C*(*t*):

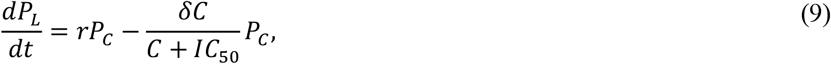

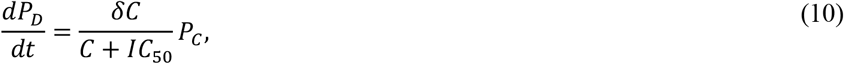

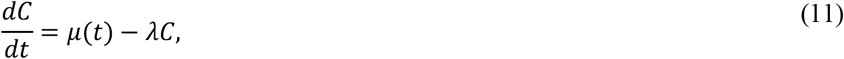

where *r* is the exponential proliferation rate of cancer cells *in vitro, δ* is the death rate of cancer cells by gemcitabine, *IC*_50_ is the drug’s half effect concentration, and λ is the decay rate of the drug (**Figure 3**C). To first determine the proliferation rate of pancreatic cancer cells *in vitro*, an exponential growth curve was fit to cell count measurements for Mia-PaCa-2 cells [77] (**Figure 3**D, parameter values **Table S2**) using simple exponential growth (i.e. setting *C*(0) = 0 in **Eq. (9)**). Fixing this growth rate and the estimate for the decay rate of drug, we then determined the antitumour efficacy of gemcitabine-loaded fibres in the cell viability experiments. Cells were treated with aliquots of simulated body fluid from gemcitabine-loaded fibres that had been incubating for 24, 48 or 72 h (**Figure 3**E). To simulate these experiments, the model is solved piecewise such that µ(*t*) = *δ*(*t* − *t*_*aliquot*_)*C*(*t*_*aliquot*_), where *t*_*aliquot*_ are the times of the drug administrations. An approximation for the concentration of drug at each time point, *C*(*t*_*aliquot*_), can be determined using the calibrated PDE model for drug release from the fibres. Fitting the drug induced death rate and *IC*_50_ gave a good approximation to the data (**Figure 3**F). The resulting parameter values from the fit of the model can be found in **Table S2**.

**Figure 3.**
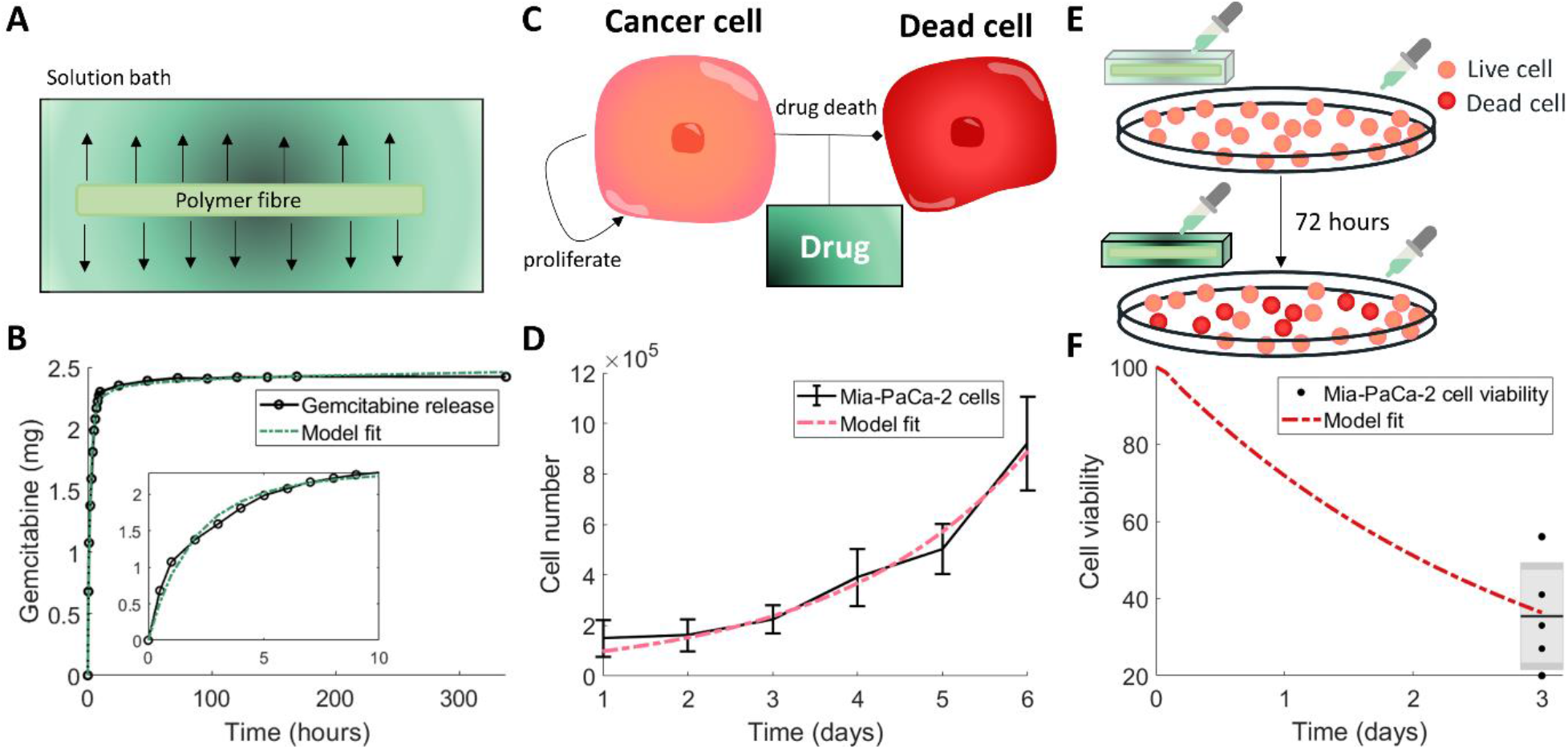
Calibration of model parameters to *in vitro* experiments. (A) Drug release profiles for gemcitabine with 3% alginate 15% PCL were measured by placing the gemcitabine-loaded alginate fibre in a solution bath and measuring the released drug concentration over time. (B) The drug concentration in the solution bath (black) was used to fit model parameters for the drug release from the fibre (green). Resulting parameters are in**Table S1**. (C) The drug-induced death rate of pancreatic cancer cells was determined by simplifying the full model assumptions to consider a homogeneous model for live cancer cells *P*_*L*_(*t*) that were proliferating and dying (become dead cells *P*_*D*_ (*t*)) through the effect of the drug gemcitabine *C*(*t*), **Eqs. (9)(11).**(D) Fitting an exponential growth curve to Mia-PaCa-2 cell proliferation *in vitro* [77] gave the growth rate of cells *r*. Values are the mean±std. (E) To measure the efficacy of the protocol, the cell viability was determined after aliquots from drug released from gemcitabine-loaded fibre were placed in a well with proliferating Mia-PaCa-2 cells at 24, 48 and 72 hours. (F) The resulting cell viability at 72 hours from the experiment depicted in (E) was used to fit the drug-induced cell death rate (**Eq. 9-11**). The data is plotted as a box and whisker plot. Resulting parameters for (D) and (F) are in **Table S2**.

### Calibration and sensitivity of pancreatic tumour growth

The VCBM simulation of pancreatic tumour growth in the absence of treatment depicts invasive and disorganised movement of cancer cells into surrounding healthy tissue (**Figure 4**A). To calibrate tumour growth parameters in the model, we used an exhaustive numerical search of the parameter space using a Latin Hypercube Sampling for *g*_*age*_, *d*_*max*_, *p*_0_ and *p*_*MCC*_, where we were minimising the least squares of the simulation with the *in vivo* tumour volume of Mia-PaCa-2 cells over 33 days (**Figure 4**B, **Table S3**). To obtain an understanding of the stochasticity in our model, we fixed the parameter values obtained and we simulated the model 100 times and plotted the tumour volume over 33 days. From **Figure 4**B, while the growth is varied at points, there are no distinct outliers or unusual tumour growth rates, and the standard deviation throughout the entire period of observation remains small. In addition, the simulations sit within the *in vivo* tumour growth measurements for pancreatic cancer growth. The histogram for the number of MCCs across the simulations (**Figure 4**C) shows only a small number of MCCs are created over the 33 days of growth, which is realistic when considering the ratio between a single cell agent in the model and a real cell in a biological tumour and which matches findings that MCCs will compose only a small subset of the tumour [78–80].

To analyse the drivers of pancreatic tumour growth dynamics in our model, we conducted a detailed sensitivity analysis. A systematic multi-parameter sensitivity analysis was performed for ***p*** = [*p*_0_, *p*_*MCC*_, *d*_*max*_, *g*_*age*_, *p*_*age*_] using weighting identified by Wells *et al*. [81] (**Figure 4**D). This sensitivity analysis can identify combinatorial influences of multiple parameters and elucidate systemic features of the model. The average tumour volume predicted by the model at day 33 for 10 simulations was recorded for each parameter set. Pairs of parameters were varied, with each cell of **Figure 4**D depicting the weighting applied to each parameter in ***p*** from 0.25, 0.75, 1.25, 1.75, and 2.25. This allowed for all combinations of alterations for two parameter values to be tested.

The time taken for a cell to prepare for mitosis, *g*_*age*_, has the greatest impact on final tumour volume (**Figure 4**D). Increasing *g*_*age*_ decreases tumour volume and conversely a decrease in *g*_*age*_ increases the final volume. As a result, the model predicts that if cells take longer to move through the cell cycle and undergo mitosis this w ill result in a smaller tumour volume. Reducing the maximum distance, a cell can be from the periphery and still proliferate, *d*_*max*_, also appears to have a decreasing effect on the final tumour volume. This is to be expected, as reducing the proliferating cell rim (through decreasing the distance from the periphery for which cells can proliferate) will reduce the number of cells available to proliferate and subsequently reduce the tumour volume. Decreasing the value of *d*_*max*_ only appears to have a significant impact on the final tumour volume when the weighting applied is ≤ 50%. In comparison with *d*_*max*_ and *g*_*age*_, the tumour volume is insensitive to changes in both the probability of a cell proliferating if it has reached mitosis, *p*_0_, and the probability of a new pancreatic cancer stem cell being created, *p*_*MCC*_. The time taken for a cell to reach adult size (when it can proliferate), *p*_*age*_, similarly has a negligible impact on the tumour volume.

**Figure 4.**
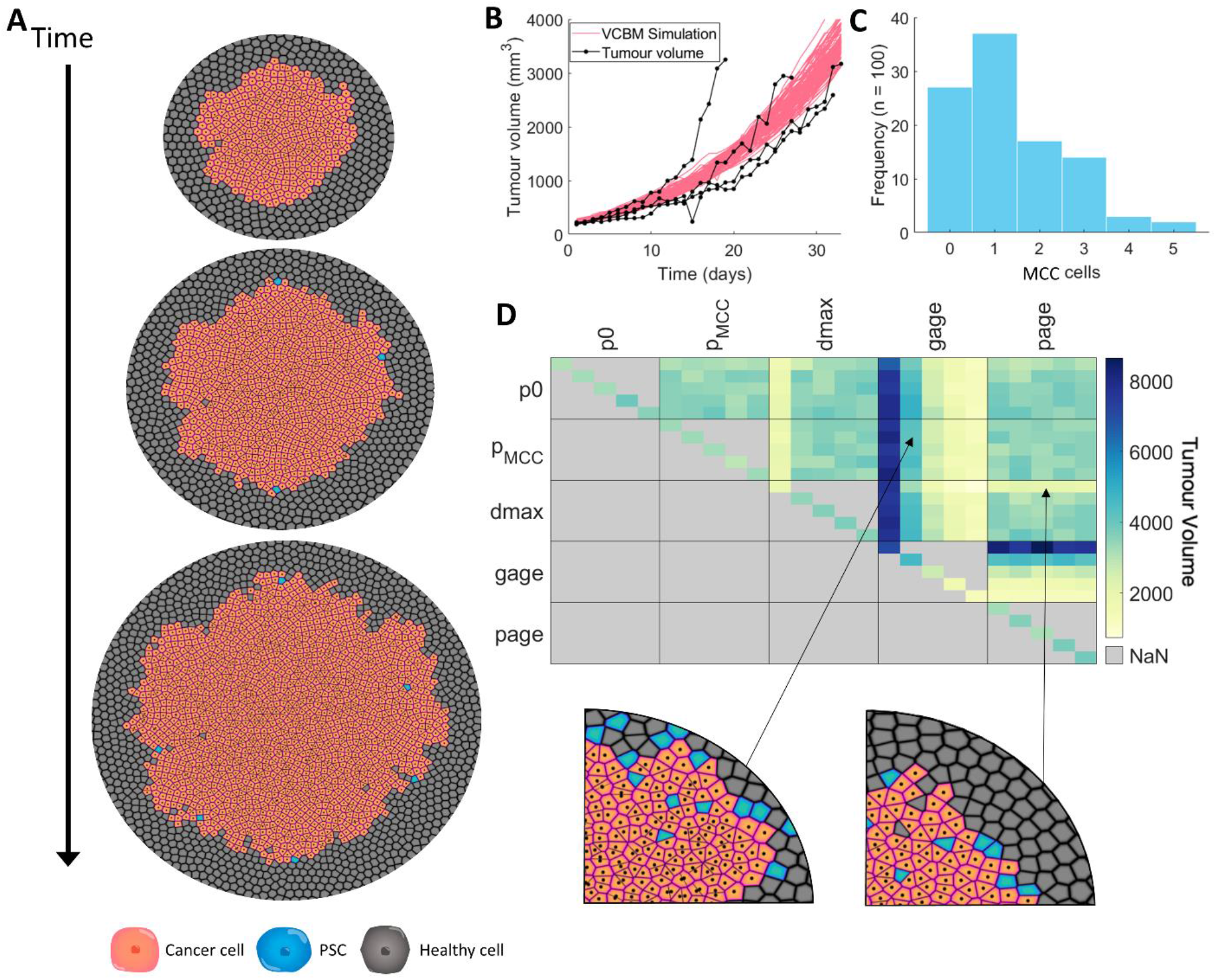
Using the VCBM to model control tumour growth. (A) Snapshots of the model simulation at 0, 5 and 10 days with cancer cells in orange, MCCs in blue and healthy cells in grey (a zoomed in version is in **Figure S6**). (B) Mia-PaCa-2 tumour volume over 33 days measured *in vivo* in mice (black, n=4). Overlaid is the tumour volume from the VCBM simulation (pink, n=100) with parameters from **Table S3**. (C) MCC counts in the VCBM simulations (n=100). (D) Sensitivity analysis of control tumour growth. Maximum tumour volume over 33 days for perturbations of parameters with weights of 0.25, 0.75, 1.25, 1.75 and 2.25, and spatial plots of large and small tumours simulated using the depicted weightings. In the heatmap, each pixel represents 30 averaged simulations with two parameters. In the boxes, the parameters vertically and horizontally in the grid are the weightings in ascending order, with each pixel being a “coordinate” representing the weighting for each parameter and the result from 30 averaged tests. Diagonal pixels only use individual parameters with different weightings.

### Intratumoural implantation provide an alternate effective protocol

Before quantifying the efficacy of gemcitabine-loaded fibres, we first looked to evaluate the impact of single point free-drug injections (**Figure 1**) of gemcitabine on the tumour volume. Simulating single point free-drug injections with the VCBM-PDE is a simplification of the full model presented in **Eqs. (1)-(4)** where *F*(*r, y, t*) = 0. More details on this can be found in the **Technical Supplementary Information**. We considered free-drug injections of gemcitabine as administered along a radial axis of the tumour in either a single dose or four free-drug injections which are rotationally symmetric (**Figure 5**A). In the case of the four injections, the total dosage is spread across the injections so that the total amount of drug administered is conserved. Simulations of the model under the different injection protocols can be found in **Figure 5**B-C and **Figure S7**. The sensitivity of parameter values governing tumour volume were again probed, now under a single administration of gemcitabine at the centre of the tumour (**Figure 5**D-E and **Figure S8**). The same trends with *g*_*age*_ and *d*_*max*_ were observed; however, an additional parameter, which represents the concentration the drug required to have an impact on the tumour volume, *IC*_50_, was found to influence the volume under further perturbations of the parameter value (**Figure 5**E). As expected, a lower value of *IC*_50_, which indicates that a smaller concentration of the drug is required for it to influence cancerous cells, leads to a lower tumour volume, while an increase leads to a higher tumour volume when compared to original estimate for *IC*_50_.

**Figure 5.**
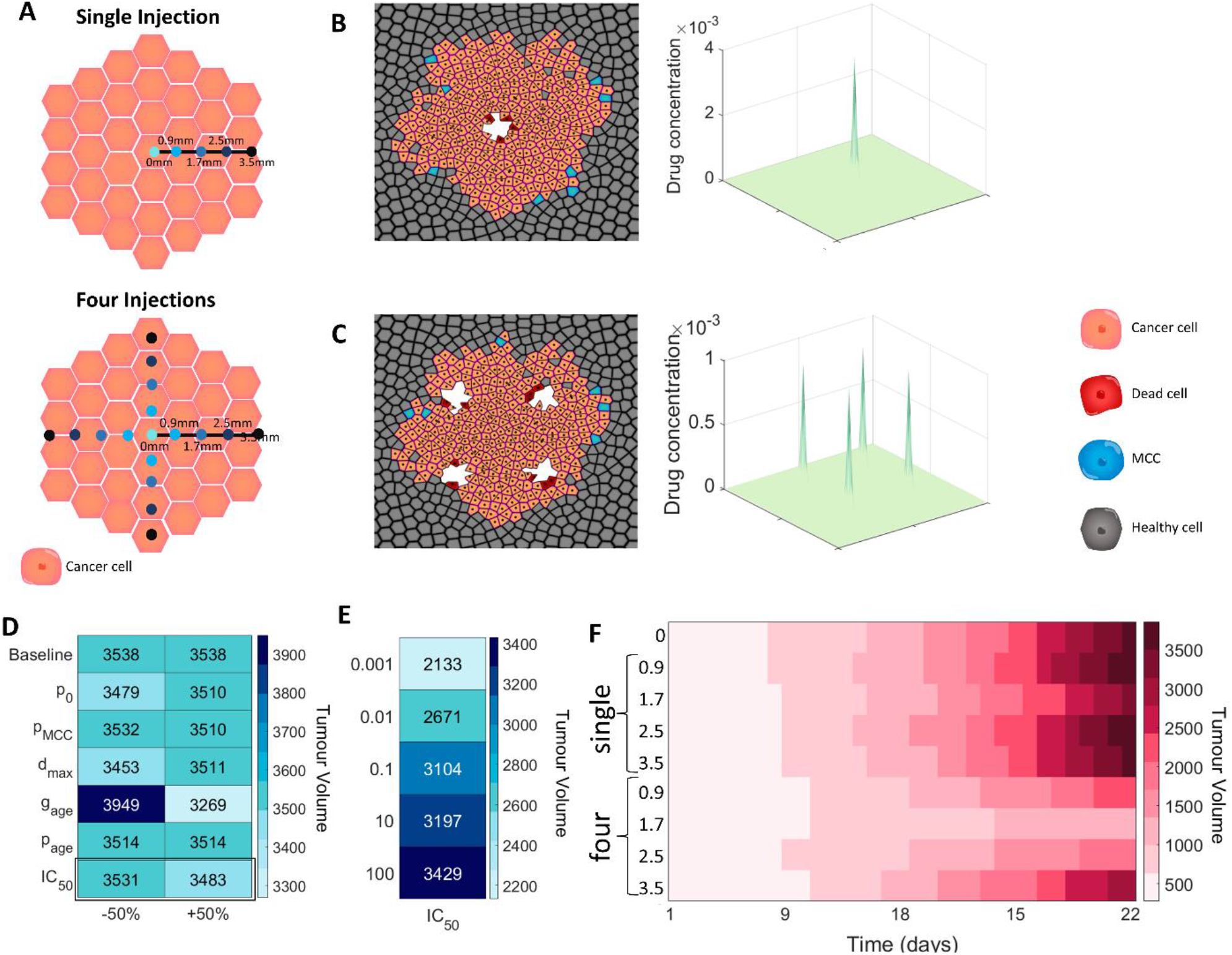
Impact of intratumoural free-drug point injections on tumour cell eradication. (A) Tumour growth was investigated under different gemcitabine single free-drug injections: central, 0.9 mm from centre, 1.7 mm from centre, 2.5 mm from centre, 3.5 mm from centre. Locations of injections on the tumour surface for single or four single free-drug injections is depicted schematically. (B) VCBM with a single central injection and the drug concentration at 24h. (C) The tumour volume with four injections placed 30µ*m* from the centre, and the drug concentration at each location at 6h. (D) Maximum tumour volume over 33 days for ±50% perturbations in parameter values compared to the normal value (i.e. baseline parameter values). (E) Maximum tumour volume over 33 days for different perturbations of *IC*_50_ compared to the normal volume. (F) The tumour volume over 33 days with each injection protocol, averaged over 10 simulations.

To determine the effect of injection placement on tumour volume over time, five placements of a single injection were considered at a distance *d*_*m*_ from the centre: a central injection (*d*_*m*_ = 0), and injections *d*_*m*_ = 0.9 mm from the centre, *d*_*m*_ = 1.7 mm from the centre, *d*_*m*_ = 2.5 mm from the centre and *d*_*m*_ = 3.5 mm from the centre (**Figure 5**A). For each of these placements, 30 simulations were run over 33 days and both the number of tumour cells and the tumour volume over time were measured (**Figure 5**F). For a single injection, distance did not impact the effectiveness of the injection and the tumour volume is qualitatively similar. There was a deviation from the consistent standard deviation width for injections further from the tumour, but this can be attributed to the method used to calculate the tumour volume in terms of how it deals with tumour structures which are not part of the central mass. The tumour volume was more significantly affected by distance in the case of four injections (**Figure 5**F), with free-drug injections further away from the centre of the tumour performing worse than those intratumoural injections. Primarily, single free-drug injections implanted peritumourally may encourage branching of external tumour structures in the model, and hence increase the calculated volume as it is based on the maximum distance from the centre of the tumour to the edge. While we present an approximation for tumour volume and placement of injections in units relevant to *in vivo* models (i.e. mm^3^ and mm respectively), more work needs to be done to validate that the efficacy of treatment predicted by the model would map to the human scale.

### Fibre location and release kinetics are a major driver of tumour arrest or tumour growth

Using the VCBM-PDE, we analysed the impact of varying the position of the fibre and the initial drug concentration on the tumour growth dynamics (**Figure 6**A). We introduced three classifications for the tumour growth dynamics: tumour eradication (i.e. a tumour volume <1mm^3^) tumour stabilisation, i.e. a tumour volume at day 33 less than the initial tumour size (≈ 100mm^3^), and tumour growth, i.e. a tumour volume on day 33 greater than the initial tumour volume. Large concentrations of gemcitabine loaded into the fibre positioned at *d*_*m*_ = 3.5 mm or *d*_*m*_ = 4.3 mm from the tumour centre were unable to stabilise or eradicate the tumour, also known as tumour arrest (**Figure 6**B-C and **Figure S10**). Once the fibre was positioned closer to the tumour centre (≤ 1.7 mm) lower concentrations of drug were sufficient to result in stabilisation of the tumour growth (**Figure 6**B). It was only with high drug concentration and centered fibres that we saw complete tumour eradication (**Figure S9**). There are large variations in the response of tumour growth to the different protocols, suggesting that tumour stabilisation or arrest might be achievable for some tumours whereas others might experience tumour growth even in the presence of drug-loaded fibre.

**Figure 6.**
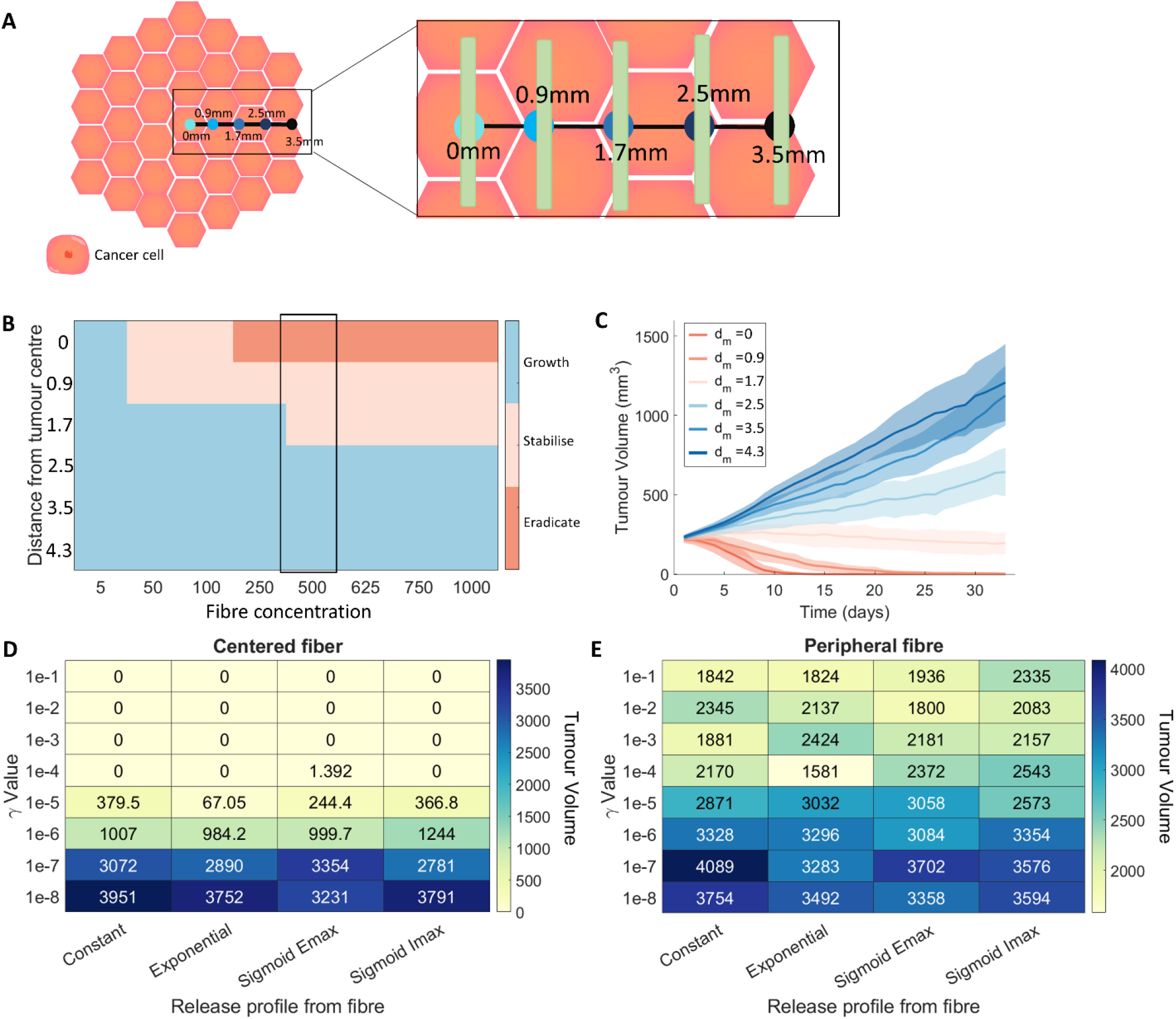
Comparison of different fibre release and placement options. (A) Tumour growth was investigated under different gemcitabine-loaded fibre placements *d*_*m*_: central, 0.9 mm from centre, 1.7 mm from centre, 2.5 mm from centre, 3.5 mm from centre and 4.3 mm from centre. Locations of fibres on tumour surface for single implantations is depicted schematically. (B) A heatmap for the averaged final state of a tumour after 33 days of simulation for different initial injection concentrations and fibre placements. “Eradicate” denotes a tumour volume below 1*mm*^3^, “stabilise” denotes a tumour volume less than the initial tumour volume, and “growth” denotes a tumour volume greater than the initial tumour volume. (C) The mean (solid lines) and standard deviation (shades areas) of the tumour volume over 33 days for different fibre placement options with corresponding values highlighted in (B). (D) The tumour volume on day 33 for different release rates (indicated by the gamma value) and release profiles with a central fibre placement. (E) The tumour volume on day 33 for different release rates (indicated by the gamma value) and release profiles with a fibre placed on the edge of the tumour (50µ*m* away from centre). See **Section TS3** of the **Technical Supplementary Information** for more details on these release functions.

To then analyse the effects of changes to the drug release profile on the tumour growth, we investigated four different release profiles: constant release, exponential release, sigmoidal Emax/Imax release profiles [82–84] (See the **Technical Supplementary Information, Section TS3**). Each of these release profiles were parameterised by a release rate γ and for the Emax and Imax curves a half-effect term η. The different release profiles were tested with the fibre placed either centrally (intratumourally) (**Figure 6**D) or on the periphery of the tumour (peritumourally) (**Figure 6**E). The four different release profiles (constant, exponential, sigmoid emax, sigmoid imax) were tested with 8 different release rates. For each parameter value,10 simulations were run over 33 days, with an initial amount of 500 *µg* of gemcitabine.

For fibres positioned in the centre (**Figure 6**D), it is possible to eradicate the tumour with all release profiles considered given a small enough value of γ. In comparison, none of the drug release profiles resulted in tumour eradication when positioned peripherally (**Figure 6**E). However, interestingly an exponential release profile with a release rate of γ = 10^−4^ results in the greatest decrease in tumour volume. This ideal release rate is likely because it allows the drug concentration to remain in the therapeutic range and kill newly developed pancreatic cancer cells aresting the process of cell proliferation. Comparing the drug release profiles (**Figure S11**), we see that the exponential release rate is similar to the sigmoid release profiles, but slightly steeper initially, suggesting that a smooth release rate with a sufficiently large initial drug release might be an optimal protocol to achieve a reduction in tumour size.

## Discussion

PDAC is a difficult-to-treat cancer with a poor prognosis. Novel therapeutic interventions are desperately needed to improve patient survival. While chemotherapy drugs, such as gemcitabine, have shown durable efficacy for pancreatic cancer, there has been little to no improvement in patient survival in the last 30 years [85]. PDACs are notorious for a dense fibrotic stroma that is interlaced with ECM [86] and is a major cause of therapeutic resistance [87]. One way of improving drug retention at the tumour site, and by consequence increase tumour eradication and patient survival, is through sustained-delivery devices (**Figure 1**). Polymeric fibres loaded with gemcitabine have shown increased therapeutic efficacy over conventional treatment delivery. To further analyse the potential of these novel therapeutic implants, we have designed a hybrid Voronoi cell-based model (VCBM)-partial differential equation (PDE) model to describe pancreatic tumour formation in healthy pancreatic tissue and the resulting effect of gemcitabine on the tumour tissue when delivered locally. With this model, we considered both the impact of a single fibre implanted with varying drug release profiles and hypothesised alternative and more effective treatment protocols.

The model was calibrated to data and with these estimates, a parameter sensitivity analysis then revealed that the fundamental driver of tumour growth in our model was the rate of cell mitosis. The idea that the cell cycling time is a fundamental part of tumour progression has been found in other mathematical models [88], suggesting that the model’s sensitivity in terms of tumour volume is in line with other models in the literature. It is also known that molecules can modulate the cell cycle of cancer cells, changing the cancer aggressivity. For example, melatonin is a hormone known for its antitumour efficacy as it significantly increases the duration of the cell cycle of human breast cancer cells [89]. Given a heterogeneous cohort of individuals with varying degrees of tumour growth rates, our model suggests that the driver of these differences is most likely the cell cycling rate. Drugs targeting this should, therefore, be considered.

Depending on the cancer type, administering an intratumoural injection of a drug can be extremely difficult and administering treatments on the periphery can be an easier course of action. Simulating the model, we found that intratumoural administration of gemcitabine-loaded fibres significantly outperforms peritumoural administration both in terms of the number of fibres and fibre placement. However, there is a threshold distance from the tumour to achieve an effective treatment, beyond which placing fibres further into the tumour bulk sees no added benefit. There is a clear benefit to increasing the dosage multiplicity and spreading the administered drug out amongst the tumour compared to a single high dose. Tumour volume was most significantly decreased when four free-drug point injections were administered compared to a single free-drug point injection. This proposes the existence of a potential threshold above which increasing the multiplicity of dosages or dosage size has a negligible effect over spreading out the dosages.

The location of the fibre and the total drug concentration in the fibre was a major driver of tumour eradication. For fibres located within the centre of the tumour with a significantly high drug concentration, it was possible to completely eradicate the tumour. Moving the fibre farther away from the centre, we found that there was no concentration of drug that would inhibit growth. This suggests that a large amount of drug from the implants is lost to the surrounding tissue, and this has detrimental effects on the efficacy of these devices. Fortunately, simulations show there is a minimal concentration of drug necessary for stabilisation, allowing these predictions to be used a way to guide dosage so that toxicity is minimised and efficacy is maximised.

The release of the drug from the fibre has a major effect on the resulting tumour volume. Implementing an exponential drug release profile, we were able to optimise the treatment to reduce the tumour size most significantly. This suggests that an initial high dosage of drug followed by a slow decline in the drug release may be an optimal protocol. This may be because it initiated a large amount of cell death initially, followed by a slower diffusion to reach remaining viable cells. While exciting, an exponential release profile needs to be tested experimentally both for its feasibility for the polymer release and to verify the predicted efficacy.

More recently, research has been focused on combining gemcitabine with other drugs to improve its efficacy. Nanoparticle albumin-bound paclitaxel (nab-paclitaxel) administered in combination with gemcitabine [9] is one of the standard of care treatment regimens that has shown an increase in overall survival in patients with advanced PDAC, as shown in a Phase I/II clinical trial [9]. A phase III clinical trial showed that gemcitabine and erlotibin also significantly increased overall survival in advanced PDAC patients compared to gemcitabine alone [90, 91].

Due to wanting to reduce the computational complexity of the VCBM, we made some simplifying assumptions that have introduced limitations into our model. To avoid simulating excessively large numbers of cells, we have chosen to scale the spatial unit appropriately so that we simulate on the order of ∼10^6^ cells. An improvement for this model, could be to parallelise the agent update step to increase the speed of the simulation. In addition, we consider only a 2-dimensional cross section of the tumour, which is a simplification given tumour’s grow in 3-dimensional environments. We feel that since we model neighbouring tissue as having a homogenous effect on tumour growth, there would be no significant impact of extending our model to 3 dimensions. Lastly, we model cell uptake by point sink terms; however, a cell would uptake drug across its surface area through drug molecule binding and internalisation. It would be possible to model this by extending the framework from a single point uptake to a uniform uptake across a cell’s defined Voronoi cell region.

There are considerable avenues for future extensions of this work, and we feel the platform we have built is easily extendable by other computational oncologists. In particular, future modelling could extend the model to account for the dense fibrotic nature of PDAC [86, 87] and investigate the impact the release and delivery of drug. In addition, the model could be used to simulate the efficacy of dual drug-loaded polymer and verify whether improvements on the current treatment protocol exist. There are many applications of degradable polymeric drug delivery systems in cancer therapy [10], for example, Rezk *et al*. [10] developed a pH-sensitive polymeric carrier to study the local delivery of anticancer drug bortezomib. They fitted the release profile of the drug from their carrier system to a mathematical formalism. Using our pancreatic cancer growth VCBM, it would be possible to feed in their drug release mechanism and simulate the efficacy under alternative protocols and predict the remaining tumour volume. Lastly, while we did not consider gemcitabine resistance in our model, it does occur in PDAC [3, 4]. A simple extension of the model could consider the impact of resistance on the performance of therapy like other works on resistance of chemotherapeutics using mathematical models [22, 92].

## Conclusion

Treatment for cancers with a poor prognosis, such as PDAC, are in vital need of novel therapeutic approaches that provide sustained, heightened, localised drug concentrations. The computational platform developed in this work can recapitulate spatially heterogeneous tumour growth and treatment with the chemotherapy drug gemcitabine. Investigating the efficacy of gemcitabine released from a degradable polymeric fibre implant, we are able to suggest that a minimum dosage for maximum efficacy exists based on the location of the device within the tumour. Furthermore, certain release profiles are significantly more effective than others, suggesting that the way in which drug is released from these devices is crucial to improving patient treatment. Moving forward, a study of this form could be used to help inform experimental design and be integrated into future device development.

## Funding

ALJ and IP were funded by the First Byte Funding Scheme from the Centre for Data Science at Queensland University of Technology. ALJ was also funded by a Fonds de recherche du Quebec – Sante International Postdoctoral Fellowship, Centre for Applied Mathematics in Biosciences and Medicine (CAMBAM). Funding from the Illawarra Cancer Carers and the PanCare Foundation (APP1165978, administered by Cancer Australia) awarded to KLV is also gratefully acknowledged. PSK was supported by the Australian Research Council Discovery Project (DP18010512).

## Notes

### Competing Interest Statement

The authors have declared no competing interest.

